# Crotamiton derivative JM03 extends lifespan and improves oxidative and hypertonic stress resistance in *Caenorhabditis elegans* via inhibiting OSM-9

**DOI:** 10.1101/2021.09.19.460995

**Authors:** Keting Bao, Jiali Feng, Wenwen Liu, Zhifan Mao, Lingyuan Bao, Tianyue Sun, Zhouzhi Song, Zelan Hu, Jian Li

**Author notes:** Correspondence: Jian Li; Zelan Hu. Keting Bao, Jiali Feng and Wenwen Liu contributed equally to this work.

## Abstract

While screening our in-house 1,072 marketed drugs for their ability to extend the lifespan using *Caenorhabditis elegans* (*C. elegans*) as an animal model, crotamiton (*N*-ethyl-o-crotonotoluidide) showed anti-aging activity and was selected for further structural optimization. After replacing the ortho-methyl of crotamiton with ortho-fluoro, crotamiton derivative JM03 was obtained and showed better activity in terms of lifespan-extension and stress resistance than crotamiton. It was further explored that JM03 extended the lifespan of *C. elegans* through osmotic avoidance abnormal-9 (OSM-9). Besides, JM03 improves the ability of nematode to resist oxidative stress and hypertonic stress through OSM-9, but not osm-9/capsaicin receptor related-2 (OCR-2). Then the inhibition of OSM-9 by JM03 reduces the aggregation of Q35 in *C. elegans* via upregulating the genes associated with proteostasis. SKN-1 signaling was also found to be activated after JM03 treatment, which might contribute to proteostasis, stress resistance and lifespan extension. In summary, this study explored a new small molecule derived from crotamiton, which has efficient anti-oxidative, anti-hypertonic and anti-aging effects, and could further lead to promising application prospects.

## Introduction

In spite the fact that aging is an inevitable process, many efforts have been made to uncover drugs which could delay aging. Numerous aging associated signaling pathways, discovered in *C. elegans*, are found to be conserved in the mammals ^1^. Moreover, many compounds, which extended the lifespan of *C. elegans*, also showed anti-aging effects in the mice model. For example, urolithin A was found to prolong the lifespan and normal activity including mobility and pharyngeal pumping in *C. elegans*, and it also improved the exercise capacity in mice with age-related decline of muscle function ^2^. Similarly, rapamycin has been also found to increase the lifespan in worms ^3^, yeast ^4^ and flies ^5^, as well as mean and maximum lifespans in mice ^6^. Metformin, a first-line drug for type 2 diabetes treatment, has been widely studied to extend the lifespan both in *C. elegans* ^7–9^ and mice ^10, 11^.

In order to discover novel anti-aging compounds, we screened our in-house 1,072 marketed drugs using *C. elegans* as an animal model for their ability to extend the lifespan. As marketed drugs generally have definite pharmacokinetics and pharmacodynamics properties and are useful for drug repurposing, our research group is focused on searching compounds for drug repurposing. Herein, in this study, the approved drug crotamiton has been found to show anti-aging activity for the first time and was further selected for the structural optimization.

Crotamiton is an inhibitor of TRPV4 (Transient Receptor Potential Vanilloid-4) channels and has been used as anti-scabies and anti-itch agent in humans for nearly 70 years ^12^. TRPV subfamily proteins are encoded by five genes in *C. elegans*, including *osm-9* (*<u>osm</u>*otic avoidance abnormal), *ocr-1* (*<u>o</u>*sm-9/*<u>c</u>*apsaicin receptor *<u>r</u>*elated), *ocr-2*, *ocr-3* and *ocr-4* ^(ref. 13)^. Only loss of *osm-9* or *ocr-2* in worms resulted in the lifespan extension ^14^. OSM-9 and OCR-2 can form heterotetrameric channels which transduce signals from olfactory, nociceptive, and serotonergic neurons ^15–17^, however the role of OSM-9 and OCR-2 in the regulation of stress resistance involves different mechanisms ^18^. It has been shown in previous studies that the inactivation of OCR-2 extends the L1 Starvation Survival, while null mutations in *osm-9* did not alter L1 starvation survival ^19^. *Osm-9* null mutants showed more resistance to oxidative or hypertonic stress than the control worms ^20^. Noticeably, it was reported that taurine, an essential amino acid involved in various physiological functions, promoted longevity of *C. elegans* in oxidative stress condition by inhibiting OSM-9 but not OCR-2^(ref. 18)^. Taken together, further extensive research is still needed to decipher the downstream signaling pathways after OSM-9 or OCR-2 activation.

With an aim to get a potential anti-ageing tool molecule, structural optimization based on crotamiton led to the identification of JM03. This molecule displayed better activity than crotamiton in terms of the lifespan extension and stress resistance of *C. elegans*. To further decipher the mechanisms of JM03 involved in the anti-aging activity, this study was conducted with special emphasis on its interaction with OSM-9 or OCR-2.

## Results

### Crotamiton prolongs lifespan of *C. elegans*

To identify candidate anti-aging compounds, we initially performed a phenotypic screening of our in-house 1,072 marketed drugs with 15 worms per concentration (100 μM) using an *C. elegans* model for their ability of lifespan extension (Figure 1-source data 1). Thereafter, 125 drugs which showed up to 10% increase in mean lifespan extension as compared to the controls were selected for the secondary screening with 30 worms per concentration (100 μM) (Figure 1-source data 2). Finally, 10 drugs which showed up to 10% increase in mean lifespan extension were chosen for the third screening with 60 worms per concentration (400 μM, 100 μM and 25 μM), respectively (Figure 1-source data 2). Apart from recently reported verapamil hydrochloride ^21^ and chlorpropamide ^22^ listed in our screening results, crotamiton was finally selected as one hit compound with significant effect on *C. elegans* lifespan extension in this study (*P* < 0.01) (Fig. 1a). The toxicity of crotamiton was evaluated by following parameters: (1) The reproductive capacity was not changed in crotamiton-treated worms at 400 μM (Fig. 1b); (2) For the normal human fetal lung fibroblasts cells MRC-5, crotamiton significantly increased its viability at 200 μM and showed no toxicity even up to 400 μM (Fig. 1c).

**Fig. 1.**
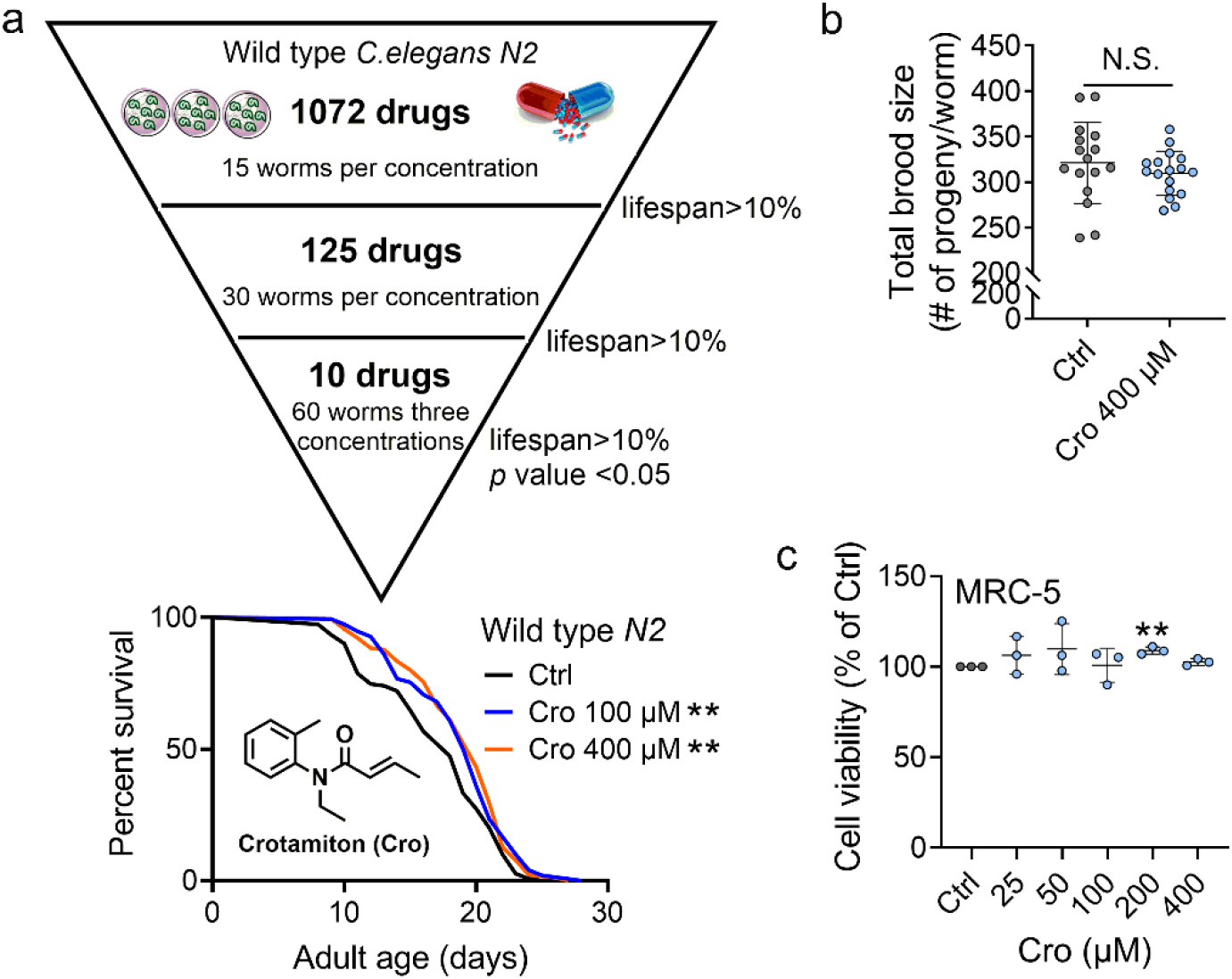
Crotamiton extends the lifespan of *C. elegans*. (**a**) Phenotypic screening led to the discovery of crotamiton as a hit compound for prolonging the lifespan in wild type (*N2*) worms. Data were compared using the Log-rank test and statistics have been mentioned in Table S1. (**b**) The total brood size of crotamiton-treated *N2* worms. Control n = 16 and crotamiton n = 17. (**c**) The viability of crotamiton-treated MRC-5 cells. *P*-values by Student t-test. *P* = 0.0015 for Cro 200 μM. (**b**, **c**) Data have been represented as the mean ± SD, and comparisons are made using Student t-test. The graphics represent a compilation of at least 3 independent experiments. * *P* < 0.05, ** *P* < 0.01.

In order to exclude the fact that the anti-aging effect of crotamiton was due to its anti-scabies activity, two anti-scabies drugs, benzyl benzoate and permethrin, were also examined along with crotamiton in the same experiment. However, both of the compounds failed to extend the worm lifespan (Fig. S1), which indicated that it was not the anti-scabies activity of crotamiton which led to the lifespan extension. Moreover, crotamiton showed no significant effect on the bacterial growth, which further ascertained that the anti-aging effect is not the result of insufficient food (Fig. S2a).

### JM03, the derivative of crotamiton, has better life extension activity in *C. elegans*

Structural optimization of crotamiton was conducted to identify more potent compounds with better anti-aging activity. The synthetic route for compounds JM01-JM15 was shown in Fig.2a. Treatment of the substituted N-ethylaniline with acryloyl chloride derivatives and potassium carbonate in dichloromethane at room temperature resulted in the formation of JM01-JM05, JM10, JM12, JM13, JM15, **9**-**11** in about 90-98% yield. Then, compounds JM10, JM12, JM15, **9**-**11** were conveniently hydrolyzed to provide the corresponding acid JM06-JM09, JM11, JM14.

Studies on the relationship between structure and activity (Fig.S3) showed the removal of the methyl group from the *ortho*-position (Crotamiton) to *meta*-position (JM01) led to the minor improvement in the activity for the R1 substituent group at benzene ring. However, moving the methyl group to the *para*-position (JM02) resulted in better activity. Replacing the *ortho*-methyl with *ortho*-fluoro (JM03), chloro (JM04) and bromo (JM05) significantly increased the activity. Additionally, incorporation of the carboxyl (JM06, JM07) on the benzene ring did not increase the activity. Since the introduction of a carboxyl at the terminal of alkenyl of crotamiton (JM08) improved the activity, we conducted additional modification based on the potent compounds JM02 and JM03. Unfortunately, no remarkable increased activity was observed after this step (Fig.S4). Moreover, the movement of the fluoro substituent from the *ortho*-position (JM03) to the *para*-position (JM13) had no effect on the activity. Further adding carboxyl (JM14) or ethoxycarbonyl (JM15) was found to be detrimental. Considering the introduction of fluorine substituents into drugs can enhance biological activity and increase chemical or metabolic stability ^23^, JM03 was selected for the following study (Fig. 2a). The lifespan of worms treated with JM03 increased significantly as compared to those treated with crotamiton (Fig. 2b). It has been reported that aging would lead to slower and uncoordinated body movement in *C. elegans* ^24^. Therefore, keeping this in view, we measured the age-dependent muscle deterioration and diminished pharyngeal pumping rate in worms to assess the healthspan of worms treated with JM03. It was found that JM03 did not change the body bend rate of *C. elegans* at different age (Fig. 2c). JM03-treated groups exhibited increased pharyngeal pumping rate at day 9 (Fig. 2d). Additionally, no changed reproductive capacity was observed in JM03-treated worms (Fig. 2e). Similar to crotamiton, JM03 showed no toxicity against MRC-5 cells and increased their viability at 200 and 400 μM (Fig. 2f). Moreover, the anti-aging effect of JM03 is not the result of insufficient food (Fig. S2b).

**Fig. 2.**
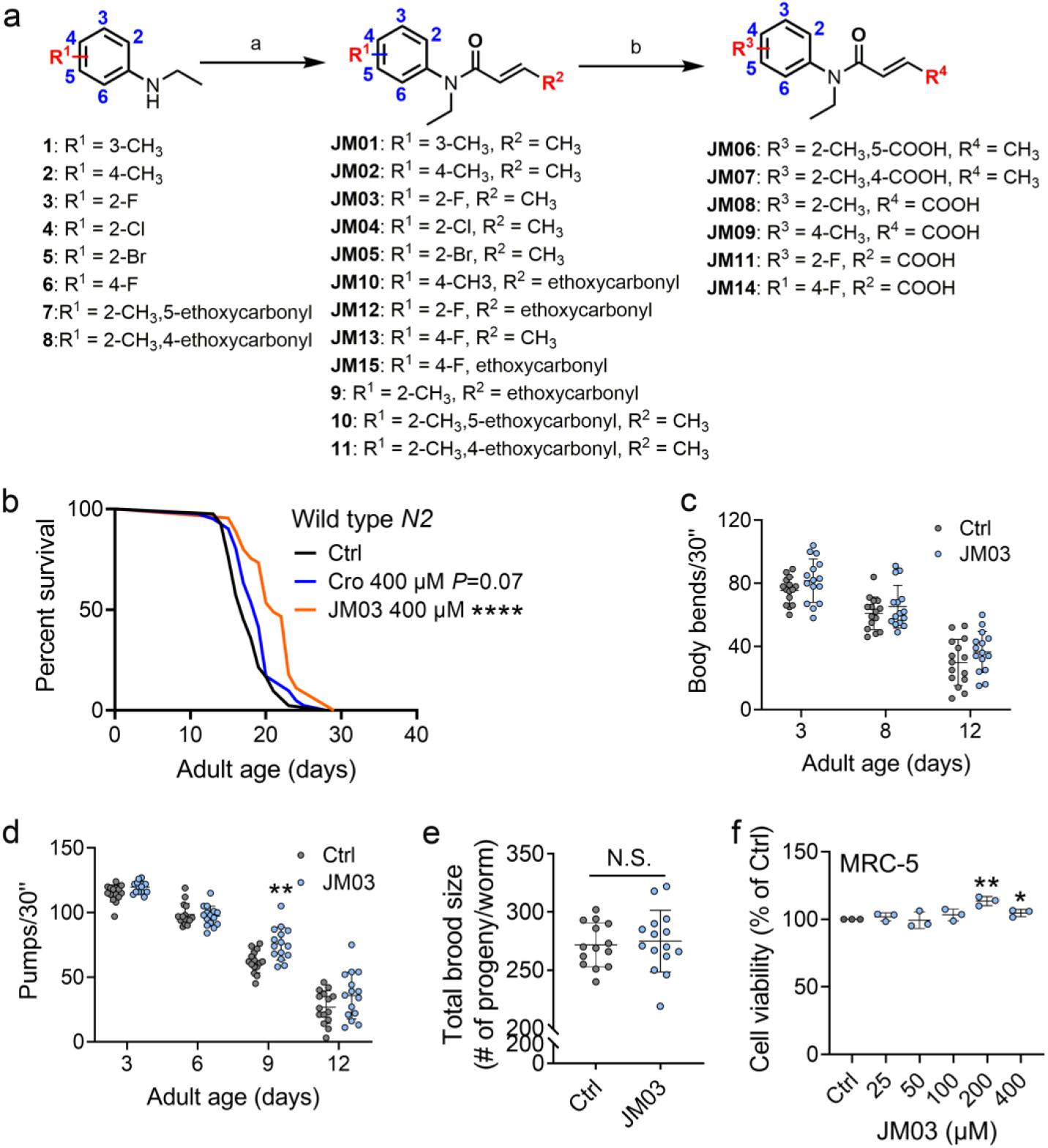
JM03 has better lifespan extension activity in *C. elegans*. (**a**) Synthesis of compounds JM01−JM15. Reagents and conditions: (a) Acryloyl chloride derivatives, K2CO3, CH2Cl2, 0 °C to rt; (b) 1M NaOH (aq.), CH3OH, rt. (**b**) JM03 prolonged lifespan in wild type (*N2*) worms. Data were compared using the Log-rank test and statistics have been mentioned in Table S1. *P* = 0.0737 for Cro 400 μM. *P* < 0.0001 for JM03 400 μM. (**c**) The mobility of JM03-treated *N2* worms by analyzing the body bend rate at day 3, 8 and 12. Control n = 15 and JM03 n = 15. (**d**) The pharyngeal pumping rate of JM03-treated *N2* worms. Control n = 15 and JM03 n = 15 at day 3, 6, 9 and 12. *P*-values by two-way AVOVA. *P* = 0.0015 for 9 days. (**e**) The total brood size of JM03-treated *N2* worms. Control n = 14 and JM03 n = 15. (**f**) The viability of JM03-treated MRC-5 cells. *P*-values by Student t-test. *P* = 0.0023 for JM03 200 μM. *P* = 0.0412 for JM03 400 μM. (**c**-**d**) Data have been represented as the mean ± SD, and comparisons are made using two-way AVOVA. (**e**-**f**) Data have been represented as the mean ± SD, and comparisons are made using Student t-test. The graphics represent a compilation of at least 3 independent experiments. * *P* < 0.05, ** *P* < 0.01, **** *P* < 0.0001.

### JM03-induced extension of lifespan depends on OSM-9 in *C. elegans*

Crotamiton is an inhibitor of TRPV4 channel ^12^, which shows similarity to the *C. elegans* channels OSM-9 (26% amino acid identity, 44% identity or conservative change) and OCR-2 (24% identity, 38% identity or conservative change) ^25^. In *C. elegans*, lacking of TRPV channel OSM-9 or OCR-2 resulted in the lifespan extension^26^. Therefore, we further performed the lifespan analysis on *osm-9* or *ocr-2* knockdown worms to investigate the mechanism of JM03. As shown in Fig. 3a and 3b, knockdown of *osm-9* or *ocr-2* via RNAi (Fig. 3d and 3e) extended the lifespan of worms compared to the empty vector group. This indicated that TRPV inhibition extended *C. elegans* lifespan. Notably, JM03 failed to extend the lifespan of osm-9 knockdown worms (Fig. 3a), but still extended the lifespan of *ocr-2* knockdown worms (Fig. 3b). Consistently, JM03 was found to be unresponsive to the lifespan of *osm-9(ky10)* mutants (Fig. 3c). These results suggested that OSM-9 not OCR-2, played a leading role in JM03-mediated longevity.

**Fig. 3.**
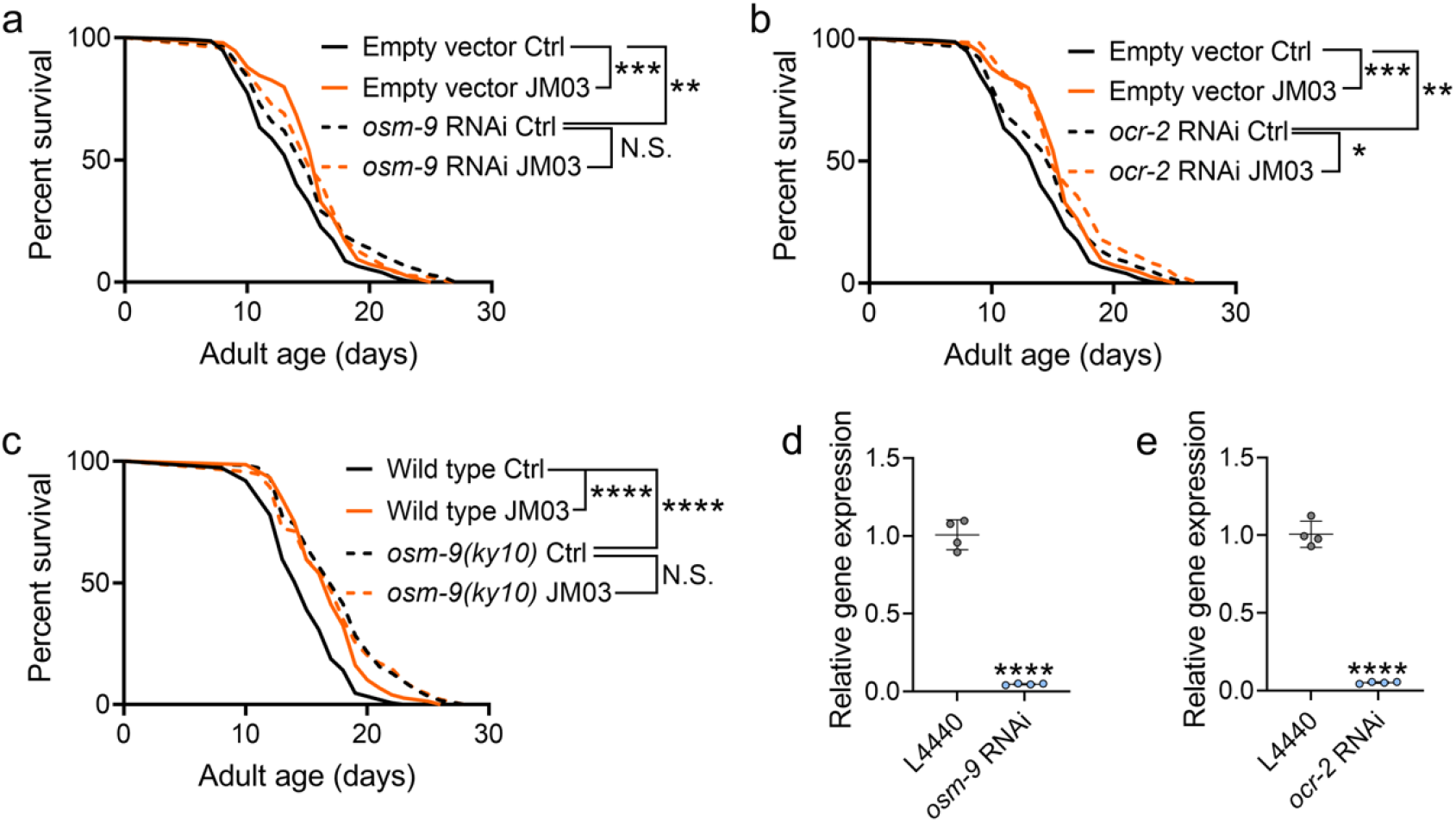
JM03-induced lifespan extension depends on OSM-9. (**a**) JM03 failed to extend the lifespan of *osm-9* RNAi worms. *P*-values by Log-rank test. *P* = 0.0002 between Empty vector Ctrl and Empty vector JM03. *P* = 0.0028 between Empty vector Ctrl and *osm-9* RNAi Ctrl. (**b**) JM03 extended the lifespan of *ocr-2* RNAi worms. *P*-values by Log-rank test. *P* = 0.0002 between Empty vector Ctrl and Empty vector JM03. *P* = 0.0084 between Empty vector Ctrl and *ocr-2* RNAi Ctrl. *P* = 0.0259 between *ocr-2* RNAi Ctrl and *ocr-2* RNAi JM03. (**c**) JM03 failed to extend the lifespan of *osm-9(ky10)* mutants. *P*-values by Log-rank test. *P* < 0.0001 for Wild type JM03 and *osm-9(ky10)* Ctrl. (**d**) The transcriptional level of *osm-9* decreased after RNAi treatment. *P*-values by Student t-test. *P* < 0.0001 for *osm-9* RNAi. (**e**) The transcriptional level of *ocr-2* decreased after RNAi treatment. *P*-values by Student t-test. *P* < 0.0001 for *ocr-2* RNAi. (**a**-**c**) Data are compared using the Log-rank test and statistics have been mentioned in Table S2 and S3. (**d**-**e**) Data have been represented as the mean ± SD, and comparisons are made using Student t-test. The graphics represent a compilation of at least 3 independent experiments. * *P* < 0.05, ** *P* < 0.01, *** *P* < 0.001, **** *P* < 0.0001.

### JM03 improves the ability of nematode to resist oxidative and hypertonic stress through OSM-9

It has been previously shown that the loss of OSM-9 enhanced the resistance of nematodes to the oxidative and hypertonic stress ^20^. Therefore, we also evaluated the efficacy of JM03 under the oxidative or hypertonic stress condition. As shown in Fig. 4a, the lifespan of *C. elegans* under paraquat-induced oxidative stress condition was significantly increased in JM03-treated group compared with control or crotamiton-treated group. Then, we examined whether the effect of JM03 under oxidative stress condition is mediated via OSM-9 and OCR-2. It was showed that JM03 treatment didn’t increase the lifespan *osm-9(ky10)* mutants (Fig. 4b), but increased the lifespan of *ocr-2(ak47)* mutants under paraquat-induced oxidative stress condition (Fig. 4c), which suggested OSM-9 is required for JM03 to improve the anti-oxidative stress ability.

**Fig. 4.**
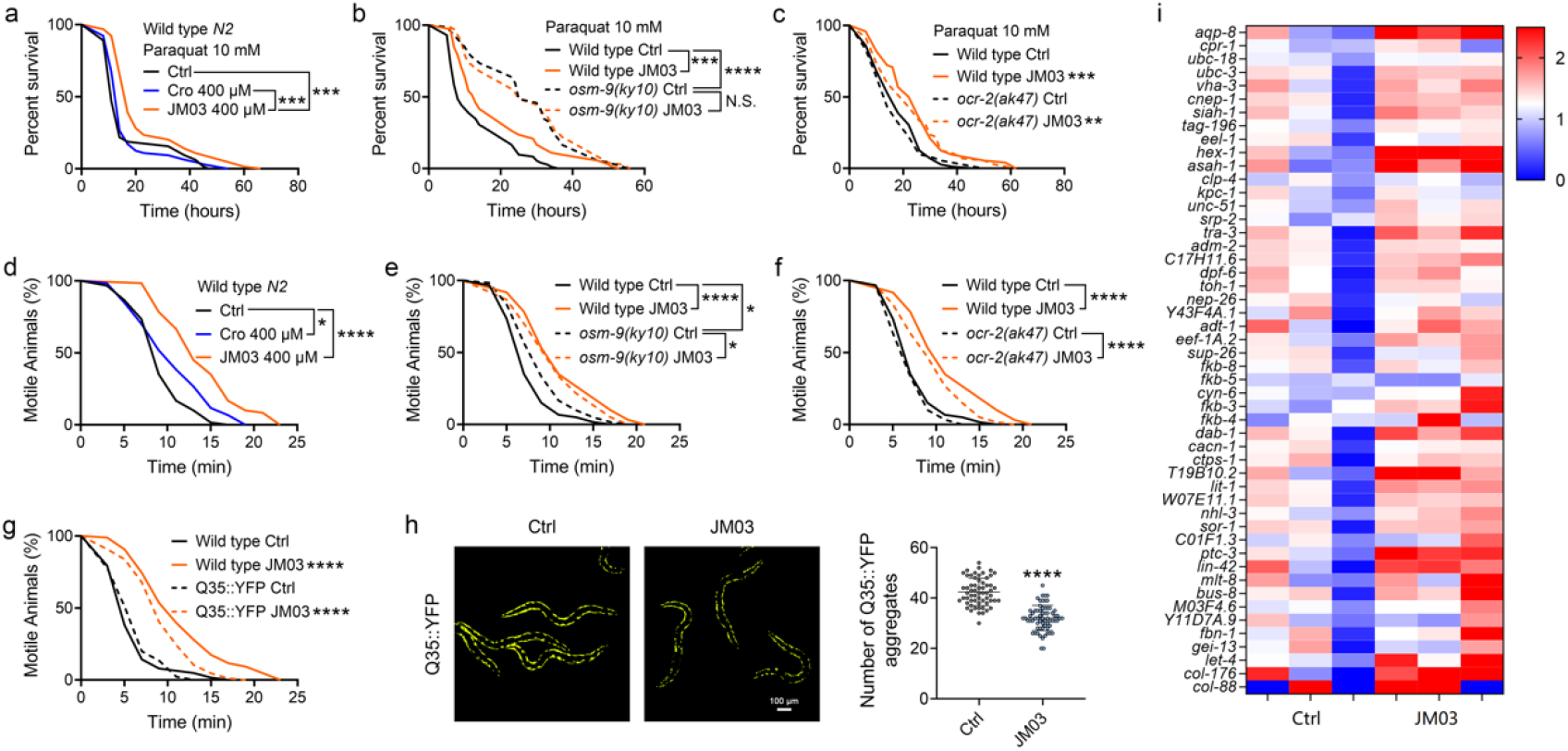
OSM-9 inhibition induced by JM03 has beneficial effect for *C. elegans* lifespan under oxidative and hypertonic stress conditions. (**a**) JM03 extended the lifespan of wild type (*N2*) worms under paraquat-induced oxidative stress condition. *P*-values by Log-rank test. *P* = 0.0001 between Ctrl and JM03 400 μM. *P* = 0.0002 between Cro 400 μM and JM03 400 μM. (**b**) JM03 failed to extend the lifespan of *osm-9(ky10)* mutants under oxidative stress condition. *P*-values by Log-rank test. *P* = 0.0009 between Wild type Ctrl and Wild type JM03. *P* < 0.0001 between Wild type Ctrl and *osm-9(ky10)* Ctrl. (**c**) JM03 extended the lifespan of *ocr-2(ak47)* mutants under oxidative stress condition. *P*-values by Log-rank test. *P* = 0.0005 between Wild type Ctrl and Wild type JM03. *P* = 0.0024 between *ocr-2(ak47)* Ctrl and *ocr-2(ak47)* JM03. (**d**) JM03 significantly reduced the paralysis for wild type worms under NaCl-induced hypertonic stress condition. *P*-values by Log-rank test. *P* = 0.0171 between Ctrl and Cro 400 μM. *P* < 0.0001 between Ctrl and JM03 400 μM. (**e**) JM03 reduced the responsiveness for *osm-9(ky10)* mutants under hypertonic stress condition. *P*-values by Log-rank test. *P* < 0.0001 between Wild type Ctrl and Wild type JM03. *P* = 0.0128 between Wild type Ctrl and *osm-9(ky10)* Ctrl. *P* = 0.0254 between *osm-9(ky10)* Ctrl and *osm-9(ky10)* JM03. (**f**) JM03 significantly reduced the paralysis for *ocr-2(ak47)* mutants similar to wild type worms under hypertonic stress condition. *P*-values by Log-rank test. *P* < 0.0001 between Wild type Ctrl and Wild type JM03. *P* < 0.0001 between *ocr-2(ak47)* Ctrl and *ocr-2(ak47)* JM03. (**g**) JM03 significantly reduced the paralysis for Q35::YFP worms similar to wild type worms under hypertonic stress condition. *P*-values by Log-rank test. *P* < 0.0001 between Wild type Ctrl and Wild type JM03. *P* < 0.0001 between Q35::YFP Ctrl and Q35::YFP JM03. (**h**) JM03 significantly reduced the Q35::YFP aggregation. *P*-values by Student t-test. *P* < 0.0001 for JM03 treatment. (**i**) Putative proteostasis genes differentially upregulated by JM03 treatment in wild type worms by transcriptome analysis. (**a**-**g**) Data are compared using the Log-rank test. (**h**) Data have been represented as the mean ± SD, and comparisons are made using Student t-test. The graphics represent a compilation of at least 3 independent experiments. * *P* < 0.05, ** *P* < 0.01, *** *P* < 0.001, **** *P* < 0.0001.

In addition, the motile time of *C. elegans* under hypertonic stress condition revealed by motility assays was significantly increased in JM03-treated group compared with control or crotamiton-treated group (Fig. 4d). We also examined whether the effect of JM03 under hypertonic stress condition is mediated via OSM-9 and OCR-2. *Osm-9(ky10)* mutants exhibited increased motility and viability upon prolonged exposures to high osmotic environments compared with wild type *N2* (Fig. 4e), while *ocr-2(ak47)* mutants exhibited motility and viability similar to wild type *N2* (Fig. 4f). For o*sm-9(ky10)* mutants, the increased significance of the motile time under hypertonic stress condition is reduced after JM03 treatment (Fig. 4e). But for *ocr-2(ak47)* mutants, JM03 still significantly increased the motile time of *C. elegans* under hypertonic stress condition (Fig. 4f). Taken together, these results suggested that the OSM-9 inhibition by JM03 increased the anti-oxidative and anti-hypertonic stress ability of *C. elegans*.

### JM03 reduces the aggregation of Q35 in *C. elegans* via upregulating the genes associated with proteostasis

It has been reported that oxidative ^27^ and hypertonic ^20^ stress enhances rapid and widespread protein aggregation and misfolding in *C. elegans*. Here, we investigated the efficacy of JM03 to reduce the aggregation of protein using Q35::YFP, a worm strain expressing polyglutamine (Q35) containing yellow fluorescent (YFP) protein in their body wall muscle. Q35::YFP is normally fully soluble in the muscles cells of young worms, but undergoes a slow, progressive aggregation as *C. elegans* ages ^28^. Here, we observed the anti-hypertonic stress ability of Q35::YFP and wild type worms was similar (Fig. 4g). JM03 significantly increased the motile time of Q35::YFP or wild type worms exposed to 500 mM NaCl, which is consistent with results shown in Fig. 4d. Meanwhile, Q35::YFP aggregation was significantly reduced when treated with JM03 (Fig. 4h), which suggested JM03 reduced the aggregation of protein in *C. elegans*.

To gain a more detailed picture of the genetic expression after JM03 treatment, worms fed with JM03 or DMSO for 10 days were processed for RNA-sequencing to analyze the altered mRNA abundance (Figure 4-source data, NCBI Gene Expression Omnibus, GSE19373). It was reported that the improved proteostasis capacity of the *osm-9* null mutant was due to altered expression of genes encoding components of the proteostasis network (including protein degradation, protein synthesis, protein folding and so on) ^20^. Interestingly, these genes that are upregulated in osm-9 null mutant and play known or presumptive roles in proteostasis, were also upregulated in JM03-treated worms (Fig 4i). Taken together, these results supported the notion that JM03 upregulates the genes associated with proteostasis through OSM-9 leading to enhanced proteostasis capacity, which may improve the ability of nematode to resist oxidative stress and hypertonic stress.

### JM03 activates the SKN-1 stress response pathway in *C. elegans*

The transcription factors DAF-16 ^(ref. 29, 30)^ and SKN-1 play important roles in regulating stress resistance, longevity and proteostasis ^31, 32^. Therefore, we examined the effect of JM03 on DAF-16 and SKN-1 pathway. JM03 prolonged the lifespan of *daf-16(mu86)* null mutant (Fig. S5), suggesting that DAF-16 is not required for JM03-induced lifespan extension. On the contrary, JM03 did not prolonged the lifespan of *skn-1(zu135)* mutants with loss of function mutation in all SKN-1 isoforms ^33^, indicating that SKN-1 played an essential role in JM03-induced positive effects (Fig. 5a). Given the dependency of the transcription factor SKN-1 in JM03-induced lifespan extension, we further examined our RNAseq dataset to determine whether expression of target genes of SKN-1 might be perturbed by JM03 treatment. We found that *skn-1* and its target genes, such as *gst-4*, *gst-6*, *gst-7*, *gcs-1*, *prdx-3* and *mtl-1* were upregulated by JM03 (Fig. 5b).

**Fig. 5.**
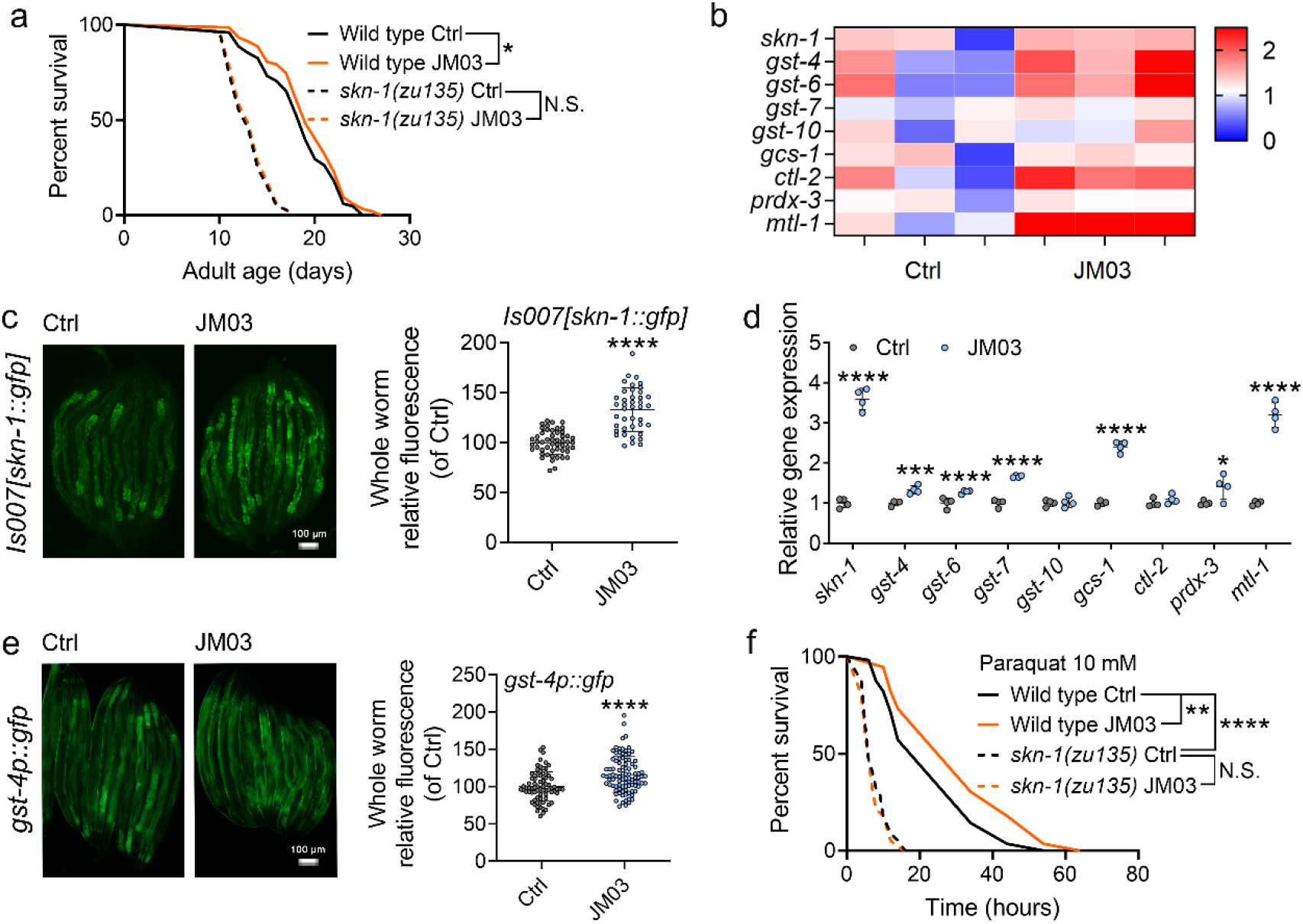
JM03-induced lifespan extension through SKN-1 pathway. (**a**) JM03 treatment failed to extend the lifespan of *skn-1(zu135)* mutants. *P*-values by Log-rank test. *P* = 0.0569 between Wild type Ctrl and Wild type JM03. (**b**) *Skn-1* and its targets genes upregulated by JM03 in wild type worms by transcriptome analysis. (**c**) JM03 significantly increased the fluorescence intensity of *skn-1::gfp*. Scale bar = 100 µm. *P*-values by Student t-test. *P* < 0.0001 for JM03 treatment. (**d**) JM03 significantly increased the transcriptional expression of *skn-1* and *skn-1* regulated genes. *P*-values by Student t-test. *P* < 0.0001 for *skn-1*. *P* = 0.0006 for *gst-4*. *P* < 0.0001 for *gst-6*. *P* < 0.0001 for *gst-7*. *P* < 0.0001 for *gcs-1*. *P* = 0.0424 for *prdx-3*. *P* < 0.0001 for *mtl-1*. (**e**) JM03 significantly upregulated the fluorescence intensity of *gst-4p::gfp*. Scale bar = 100 µm. *P*-values by Student t-test. *P* < 0.0001 for JM03 treatment. (**f**) JM03 treatment failed to extend the lifespan of *osm-9(ky10)* and *skn-1(zu135)* mutants under oxidative stress condition. *P*-values by Log-rank test. *P* = 0.0059 between Wild type Ctrl and Wild type JM03. *P* < 0.0001 between Wild type Ctrl and *skn-1(zu135)* Ctrl. (**a**, **f**) Data are compared using the Log-rank test. The graphics represent a compilation of at least 3 independent experiments. (**c**-**e**) Data have been represented as the mean ± SD, and comparisons are made using Student t-test. * *P* < 0.05, ** *P* < 0.01, *** *P* < 0.001, **** *P* < 0.0001.

Next, we examined the effect of JM03 on the activation of the SKN-1 stress response pathway using a previously described GFP translational reporter fused to the *skn-1* promoter ^33, 34^. JM03 treatment significantly increased the intensity of GFP fluorescence driven by the native *skn-1* promoter (*Is007[skn-1::gfp]*) ^33^ (Fig. 5c). Concurrently, it also significantly increased the transcriptional expression of *skn-1* itself and *skn-1* regulated genes *gst-4*, *gst-6*, *gst-7*, *gcs-1*, *ctl-2*, *prdx-3* or *mtl-1* (Fig. 5d). Subsequently, we also confirmed the increased expression of glutathione S-transferase-4 (*gst-4*), a key downstream target of SKN-1 ^(ref. 35)^, based on the GFP fluorescence signal of *gst-4::gfp* worms (Fig. 5e). In addition, JM03 did not extend the lifespan of *skn-1(zu135)* mutants under oxidative stress condition (Fig. 5f). In conclusion, JM03 prolongs the lifespan and improves stress-resistance ability of *C. elegans* through SKN-1 pathway.

## Discussion

Drug repurposing has emerged as an effective approach for the rapid identification and development of pharmaceutical molecules with novel activities against various diseases based on the already known marketed drugs ^36^. Herein, we explored the possibility of identifying the potent drugs which could increase the longevity by screening our in-house marketed drugs. Based on the screening of 1,072 marketed drugs using lifespan extension assays in *C. elegans*, crotamiton, which was known as an anti-scabies and anti-itch agent was identified for its property to increase the lifespan. We also proved that the lifespan extension effect of crotamiton was not the result of its anti-scabies activity (Fig. S1) or the change in the nutritional value of the bacteria (Fig. S2). Thereafter, structural optimization of crotamiton led to the identification of a more potential compound JM03, which was found to show better lifespan expansion and stress resistance activity than crotamiton. Various aspects of the mechanistic action of this molecule were further explored in this study.

It has been reported that crotamiton is an inhibitor of human TRPV4 channel ^12^, which is homologous to OSM-9 and OCR-2 channels in *C. elegans* ^13^. Loss of OCR-2 or OSM-9, can result in the lifespan extension in *C. elegans* ^26^. In our study, JM03 further increased the lifespan for the *ocr-2* knockdown *C. elegans*, but was ineffective for the knockdown or knockout of *osm-9* (Fig. 3), which suggested JM03 selectively acted on OSM-9, not OCR-2. Furthermore, JM03 improved the antioxidant and anti-hyperosmotic stress resistance of wild type worms (Fig. 4a, d). Interestingly, *osm-9* mutants showed enhanced ability to resist oxidative stress and hypertonic stress (Fig. 4b, e), while *ocr-2* mutants didn’t (Fig. 4c, f). These results also supported that JM03 selectively acted on OSM-9, not OCR-2. Consistently, JM03 still had significant anti-oxidant and anti-hyperosmotic effects on *ocr-2* mutants (Fig. 4c, f), but not *osm-9* mutants (Fig. 4b, e). It is noted that OSM-9 is not the only mechanism that mediate the anti-hyperosmotic effect of JM03 because JM03 retained a slight effect on osmotic pressure resistance of the *osm-9* mutants.

OSM-9 plays major roles in transduction and regulation of signals in several sensory neurons and is important for processes such as volatile chemotaxis and osmotic avoidance ^37^. *Osm-9* null mutant was reported to show enhanced survival in hypertonic environments, not due to altered systemic volume regulation or glycerol accumulation and instead may be due to enhanced proteostasis capacity ^20^. Consistently, JM03 treatment also enhanced proteostasis capacity in *C. elegans* revealed by reduced aggregation of Q35 (Fig. 4h). Besides, the genes associated with proteostasis, that are upregulated in *osm-9* null mutant, were also upregulated in JM03-treated worms revealed by transcriptome analysis (Fig. 4i). Among these genes, the increased expression of *aquaporin-8* (*aqp-8*) in *osm-9(ok1677)* mutant was also reported by a Germany lab ^38^. Considering the essential roles of AQP-8 in sustaining the salt/water balance in various cells types and tissues, the loss/inhibition of *osm-9* might help to maintain the salt/water balance to promote proteostasis during the response to hyperosmotic stress.

To investigate which downstream signaling was activated after OSM-9 inhibition by JM03, two important stress response transcription factors DAF-16 and SKN-1 were examined ^27, 39, 40^. Lifespan analysis showed that JM03 extended the lifespan through SKN-1 (Fig. 5a), but not DAF-16 (Fig. S5). Then our RNA-sequencing and qPCR data both showed the transcriptional expression of *skn-1* itself and *skn-1* regulated genes were significantly increased in JM03-treated worms (Fig. 5b, d). The expression of SKN-1 and GST-4 was confirmed by using GFP translational reporter worms (*skn-1::gfp* and *gst-4::gfp*) (Fig. 5c, e). These results provide the evidence that JM03 activates SKN-1 signaling that regulates longevity, stress resistance and proteostasis. But how JM03 activates SKN-1 signaling after inhibiting OSM-9 remained to be studied.

In conclusion, overall results showed that JM03 increased the lifespan of *C. elegans* by inhibiting OSM-9, and then activated SKN-1, which improved proteostasis, stress resistance and lifespan extension in *C. elegans* (Fig. 6). Since OSM-9 is the homologous to mammal TRPV channels, it’s very interesting to examine whether JM03 acts selectively on a certain TRPV subtypes in mice in future studies.

**Fig. 6.**
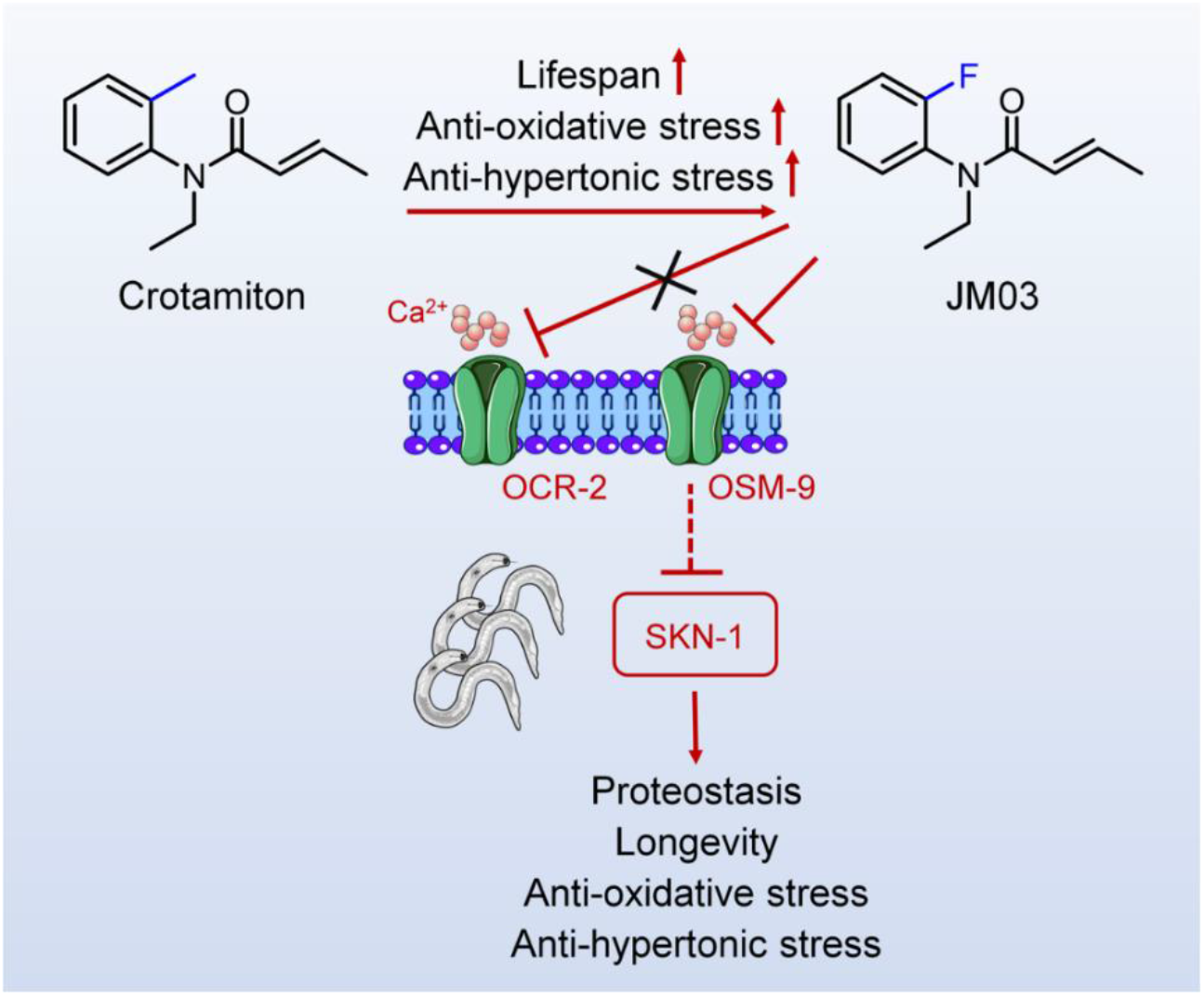
Schematic diagram of the mechanism of action of JM03 for regulating the lifespan, anti-oxidative and anti-hypertonic stress ability in *C. elegans*.

## Materials and methods

### Strains

*C. elegans* were kept at 20°C on the nematode growth media (NGM) with plates seeded with *E. coli* OP50. Strains used in this study were obtained from Caenorhabditis Genetics Center. The *C. elegans* strains used in this study were as follows: wild type Bristol strain *N2*, CX4544 *ocr-2(ak47) IV*, CX10 *osm-9(ky10) IV*, AM140 rmIs132 [unc-54p::Q35::YFP], LG333 *skn-1(zu135) (IV)/nT1[qIs51] (IV;V); ldIs7 [skn-1b/c::GFP]*, CF1038 *daf-16(mu86) (I)*, CL2166 *dvIs19 [gst-4p::GFP::NLS] (III)*, EU31 *skn-1(zu135) (IV)/ nT1[unc-?(n754);let-?] (IV;V)*.

### Lifespan analysis

Worms were synchronized with bleaching buffer, followed by the starvation in M9 buffer at the L1 stage for 24 hours. Worms were thereafter transferred to the NGM plates containing the respective compounds at L4 stage. To avoid progeny hatching, 50 μg/mL of 5-Fluorodeoxyuridine (FUdR) was added to the agar plates from day 0 to day 10. From the 10^th^ day of adulthood, all the groups were transferred to the plates without compounds or FUdR treatment until the end of life. During adulthood, worms were counted every day and transferred to the fresh plates every two days. Death was indicated by total cessation of movement in response to gentle mechanical stimulation. The survival curves were generated using GraphPad Prism 8.3.0. The log-rank (Mantel-Cox) test was used to assess the curve significance.

For lifespan screening experiments, 15 worms were cultured on each Petri dish (60 mm in diameter) containing NGM plate ^41, 42^. 1 Petri dishes in the 1^st^ round screen, 2 Petri dishes in the 2^nd^ round screen, 4 Petri dishes in the 3^rd^ round screen and 8 Petri dishes in the validation of the effect of drugs were used. In lifespan assay of mutant or control worms, more than 100 worms were used for 1 experiment and a least 3 independent experiments were performed for biological replication.

### RNAi experiment

*E. coli* strain HT115 was used for this assay. The clones used were *osm-9* (B0212.5) and *ocr-2* (T09A12.3). L4440 was used as the vector. Worms fed with the bacteria expressing L4440 or engineered to produce a gene RNAi effect were cultured until the F4 stage. One subset of the worms was confirmed to exhibit the decreased expression of the gene via qPCR. Then, the other subset of worms was synchronized for a lifespan assay on the control and JM03 400 μM treated NGM plates seeded with bacteria either expressing L4440 or engineered to produce a gene RNAi effect.

### Bacterial growth assay

A single colony of bacteria was inoculated in the LB media and cultured at 37°C. For plate assay, 30 μL of bacterial culture (OD_600_=0.12) was transferred to an NGM plate either with or without crotamiton or JM03 at a concentration of 400 μM, and cultured at 20°C. The bacteria were washed off using 1 mL M9 buffer and OD_600_ was measured every 12 h, with M9 buffer as the blank control. OD was assessed using a Hitachi U-2910 spectrometer with a 10-mm quartz cuvette. At least 3 technical and 3 biological independent replicates were performed.

### Thrashing assay

Wild type worms *N2* were transferred to the NGM plates at L4 stage and incubated with JM03 at the concentration of 400 μM. For the control and JM03 treatment groups, thrashes were counted on days 3, 8, and 12. Any change in the midbody bending direction was referred to as a thrash. Worms were placed in M9 buffer drop on an NGM plate without OP50 and allowed to adapt for 30 s. Then, the number of thrashes over 30 s were counted.

In thrashing assay, pharyngeal pumping assay and reproductive lifespan assay, more than 15 worms were used for 1 experiment and a least 3 independent experiments were performed for biological replication.

### Pharyngeal pumping assay

Wild type worms *N2* were transferred to the NGM plates at L4 stage with JM03. On days 3, 6, 9, and 12, worms were evaluated by quantifying the contractions of the pharynx over a period of 30 s.

### Reproductive lifespan assay

Wild type worms *N2* were transferred to 3.5 cm NGM plates individually at L4 stage with or without JM03 at a concentration of 400 μM. They were further moved to a fresh plate each day until 3 consecutive days without the progeny production. After transferring, plates were checked for progeny after 2 days. For each individual, the last day of the live progeny production was determined as the day of reproductive cessation.

### Cell culture and viability assay

MRC-5 cells were maintained in MEM medium (Gibco) supplemented with 10% FBS (42F6590K, Gibco), 1% non-essential amino acid solution (BI), 1% sodium pyruvate solution (BI) and 1% Penicillin-Streptomycin (100×) (Yeasen). Thereafter, the cells were cultured at 37°C in an incubator with humidified atmosphere and with 5% CO_2_. Cells were periodically checked to be mycoplasma-free using GMyc-PCR Mycoplasma Test Kit (40601ES10, Yeasen, Shanghai, China).

For the cell counting kit-8 (CCK-8) assay, MRC-5 cells (1×10^4^ cells/well) were seeded in the 96-well culture plates (100 μL/well) for 12 h. Further, different concentrations of test compounds were added to the plates in 100 μL of fresh medium (the total volume was 200 μL, DMSO < 0.1%) and incubated for 72 h. After removal of the cell culture medium, a 10% CCK-8 solution (Targetmol, C0005) in medium was added and re-incubated for 2 h. Then, the absorbance at 450 nm was measured in a microplate reader (Biotek, Vermont, USA). At least 3 technical and 3 biological independent replicates were performed.

### Hypertonic stress resistance assay

Worms were transferred to the NGM plates at L4 stage and incubated with JM03 at a concentration of 400 μM for 4 days. Approximately 60 worms were transferred to NGM plates with 500 mM NaCl and their movement (# moving/total) was assessed. Worms were defined as paralyzed if they failed to move forward upon the tail prodding. Survival was measured every two minutes until all the worms were paralyzed. More than 100 worms were used for 1 experiment and a least 3 independent experiments were performed for biological replication.

### Oxidative stress resistance assay

Worms were transferred to the NGM plates at L4 stage and incubated with JM03 at a concentration of 400 μM for 4 days. Worms were further transferred to the 24-well plate (6 worms/well) and incubated in M9 buffer containing 10 mM paraquat. Survival was measured every two hours until all the worms were died. More than 70 worms were used for 1 experiment and a least 3 independent experiments were performed for biological replication.

### PolyQ aggregation assay

PolyQ aggregation assay was performed using AM140 *C. elegans* expressing polyQ35::YFP fusion protein in muscle cells. Worms were transferred to the NGM plates at L4 stage and incubated with JM03 at a concentration of 400 μM for 4 days. Thereafter, YFP images were taken using a fluorescence microscopy (Nikon Eclipse Ti2) and the polyQ35::YFP aggregates in worms were quantified manually using imageJ software. More than 50 worms were used for 1 experiment and a least 3 independent experiments were performed for biological replication.

### Transcriptome analysis by RNA-sequencing

The transcriptome analysis by RNA sequencing was performed according to a previously published method ^43^. Wild type worms *N2* were transferred to the NGM plates at the L4 stage and incubated with JM03 at a concentration of 400 μM for 10 days. At the 10^th^ day of adulthood, worms were collected. Total RNAs were extracted using Trizol Reagent (R0016, Beyotime, Shanghai, China). Further assay and analysis were assisted by Majorbio Bio-Pharm Technology Co. Ltd (Shanghai, China) and are shown in Figure 4-source data.

### qRT-PCR analysis

Wild type worms *N2* were transferred to the NGM plates at the L4 stage and incubated with JM03 at a concentration of 400 μM for 4 days. Total RNA was extracted from *C. elegans* with a Total RNA Kit II (R6934-01, Omega, USA) and reverse transcribed using Hifair Ⅱ 1st Strand cDNA Synthesis SuperMix for qPCR (11123ES60, Yeasen, Shanghai, China) in accordance with the manufacturer’s instructions. qPCR was performed using Hieff qPCR SYBR® Green Master Mix (11201ES08, Yeasen, Shanghai, China) on a CFX96 quantitative PCR system (Bio-Rad, USA). Data were processed using CFX Maestro 1.0. The primers used are listed as follows: *ama-1*, forward: TGGAACTCTGGAGTCACACC; reverse: CATCCTCCTTCATTGAACGG. *act-1*, forward: ATGTGTGACGACGAGGTTGC; reverse: ACTTGCGGTGAACGATGGATG. *skn-1*, forward: ACAGTGCTTCTCTTCGGTAGC; reverse: GAGACCCATTGGACGGTTGA. *gst-4*, forward: TGCTCAATGTGCCTTACGAG; reverse: AGTTTTTCCAGCGAGTCCAA. *gst-6*, forward: TTTGGCAGTTGTTGAGGAG; reverse: TGGGTAATCTGGACGGTTTG. *gst-7*, forward: AGGACAACAGAATCCCAAAGG; reverse: AGCAAATCCCATCTTCACCAT. *gst-10*, forward: GTCTACCACGTTTTGGATGC; reverse: ACTTTGTCGGCCTTTCTCTT. *gcs-1*, forward: AATCGATTCCTTTGGAGACC; reverse: ATGTTTGCCTCGACAATGTT. *ctl-1*, forward: GCGGATACCGTACTCGTGAT; reverse: GTGGCTGCTCGTAGTTGTGA. *prdx-3*, forward: CTTGACTTCACCTTTGTATGCC; reverse: GGCGATCTTCTTGTTGAAATCA. *mtl-1*, forward: CAAGTGTGACTGCAAAAACAAG; reverse: GCAGTACTTCTCACAACACTTG. *osm-9*, forward: GACCGCGTAGGAGTACATGG; reverse: GAGAGGTGTGGAAGGCGAAA. *ocr-2*, forward: GCCAGTCAGCTTACCAACAC; reverse: GGTGCAGAATTTGGCGAACG.

### SKN-1 and GST-4 expression determination

CL2166 (*gst-4p*::GFP) and LG333 (*skn-1b/c*::GFP) transgenic strains were transferred to the NGM plates at L4 stage and incubated with 400 μM JM03 for 4 days. The SKN-1 and GST-4 expression was determined by the fluorescence microscopy (Nikon Eclipse Ti2). The fluorescence intensity was quantified using the ImageJ software. More than 50 worms were used for 1 experiment and a least 3 independent experiments were performed for biological replication.

### Statistical analysis

All the data are represented as mean ± SD. Statistical analysis was conducted using Graphpad Prism 8.3.0 and significant differences within treatments were determined by Log-rank (Mantel-Cox) test, two-way ANOVA or Student’s t-test. *P* ≤ 0.05 was considered statistically significant.

### General information of Crotamiton derivatives

All the reagents were purchased from commercial corporation without further purification. Nuclear magnetic resonance (NMR) spectroscopy was recorded on 400 MHz or 600 MHz Bruker spectrometer at 303 K and referenced to TMS. Chemical shifts were reported in parts per million (ppm, *δ*). High-resolution mass spectra (HRMS) data were given by Waters LCT or Agilent 6545 Q-TOF. The flash column chromatography was conducted on silica gel (200− 300 mesh) and visualized under UV light at 254 and 365 nm.

*General procedures for the synthesis of **JM01-JM05, JM10, JM12, JM13, JM15, 9-11***

To a solution of **1-8** (4.0 mmol) in dichloromethane was added acryloyl chloride derivatives (4.0 mmol) and potassium carbonate (1.66 g, 12.0 mmol) at 0 °C. Then the mixture was stirred at room temperature for about 1 h. After removing the solvent under reduced pressure, the residue was dissolved in ethyl acetate, washed with water and brine. Then the organic phase was dried with sodium sulfate and concentrated *in vacuo*. The crude compound was purified by silica gel column chromatography.

*(E)-N-ethyl-N-(m-tolyl)but-2-enamide* (**JM01**)

780 mg, 96.0 % yield. **^1^H NMR** (600 MHz, CDCl_3_) *δ* 7.33-7.27 (m, 1H), 7.16 (d, *J* = 7.5 Hz, 1H), 6.97 (s, 1H), 6.96 – 6.87 (m, 2H), 5.68 (d, *J* = 14.8 Hz, 1H), 3.80 (q, *J* = 6.9 Hz, 2H), 2.39 (s, 3H), 1.72 (d, *J* = 6.7 Hz, 3H), 1.13 (t, *J* = 7.0 Hz, 3H). ESI-HRMS [M+H]^+^ calcd for C_13_H_18_NO: 204.1383, found: 204.1389.

*(E)-N-ethyl-N-(p-tolyl)but-2-enamide* (**JM02**)

789 mg, 97.0 % yield. **^1^H NMR** (600 MHz, CDCl_3_) *δ* 7.21 (d, *J* = 7.5 Hz, 2H), 7.03 (d, *J* = 7.4 Hz, 2H), 6.94 – 6.86 (m, 1H), 5.68 (d, *J* = 15.0 Hz, 1H), 3.79 (q, *J* = 7.0 Hz, 2H), 2.39 (s, 3H), 1.71 (d, *J* = 6.7 Hz, 3H), 1.12 (t, *J* = 7.0 Hz, 3H). ESI-HRMS [M+H]^+^ calcd for C_13_H_18_NO: 204.1383, found: 204.1386.

*(E)-N-ethyl-N-(2-fluorophenyl)but-2-enamide* (**JM03**)

791 mg, 95.4 % yield. **^1^H NMR** (600 MHz, CDCl_3_) *δ* 7.39 – 7.34 (m, 1H), 7.23 – 7.16 (m, 3H), 6.99-6.91 (m, 1H), 5.63 (d, *J* = 15.0 Hz, 1H), 3.87-3.81 (m, 1H), 3.78 – 3.69 (m, 1H), 1.73 (dd, *J* = 6.9, 1.5 Hz, 3H), 1.12 (t, *J* = 7.2 Hz, 3H). ESI-HRMS [M+H]^+^ calcd for C_12_H_15_FNO: 208.1132, found: 208.1135.

*(E)-N-(2-chlorophenyl)-N-ethylbut-2-enamide* (**JM04**)

828 mg, 92.5 % yield. **^1^H NMR** (600 MHz, CDCl_3_) *δ* 7.54 – 7.50 (m, 1H), 7.37 – 7.32 (m, 2H), 7.24 – 7.20 (m, 1H), 6.97-6.91 (dq, *J* = 13.9, 6.9 Hz, 1H), 5.53 – 5.46 (m, 1H), 4.07 (dq, *J* = 14.3, 7.2 Hz, 1H), 3.48 (dq, *J* = 14.3, 7.2 Hz, 1H), 1.72 (dd, *J* = 6.9, 1.6 Hz, 3H), 1.14 (t, *J* = 7.2 Hz, 3H). ESI-HRMS [M+H]^+^ calcd for C_12_H_15_ClNO: 224.0837, found: 224.0804.

*(E)-N-(2-bromophenyl)-N-ethylbut-2-enamide* (**JM05**)

1009 mg, 94.1 % yield. **^1^H NMR** (600 MHz, CDCl_3_) *δ* 7.70 (dd, *J* = 8.0, 1.3 Hz, 1H), 7.39 (td, *J* = 7.6, 1.3 Hz, 1H), 7.28 – 7.25 (m, 1H), 7.22 (dd, *J* = 7.8, 1.6 Hz, 1H), 6.95 (dq, *J* = 13.9, 6.9 Hz, 1H), 5.48 (dd, *J* = 15.0, 1.6 Hz, 1H), 4.14 (dq, *J* = 14.3, 7.2 Hz, 1H), 3.39 (dq, *J* = 14.2, 7.2 Hz, 1H), 1.72 (dd, *J* = 6.9, 1.6 Hz, 3H), 1.15 (t, *J* = 7.2 Hz, 3H). ESI-HRMS [M+H]^+^ calcd for C_12_H_15_BrNO: 268.0332, found: 268.0308.

*Ethyl (E)-4-(ethyl(p-tolyl)amino)-4-oxobut-2-enoate* (**JM10**)

951 mg, 91.0 % yield. **^1^H NMR** (600 MHz, CDCl_3_) *δ* 7.22 (d, *J* = 8.0 Hz, 2H), 7.02 (d, *J* = 8.1 Hz, 2H), 6.82 (q, *J* = 15.3 Hz, 2H), 4.16 (q, *J* = 7.1 Hz, 2H), 3.83 (q, *J* = 7.1 Hz, 2H), 2.39 (s, 3H), 1.24 (t, *J* = 7.1 Hz, 3H), 1.15 (t, *J* = 7.1 Hz, 3H). **^13^C NMR** (150 MHz, CDCl_3_) *δ* 165.81, 163.59, 138.29, 138.26, 134.83, 130.71, 130.42, 127.84, 60.92, 44.66, 21.10, 14.10, 12.81. ESI-HRMS [M+H]^+^ calcd for C_15_H_20_NO_3_: 262.1438, found: 262.1435.

*Ethyl (E)-4-(ethyl(2-fluorophenyl)amino)-4-oxobut-2-enoate* (**JM12**)

960 mg, 90.5 % yield. **^1^H NMR** (600 MHz, CDCl_3_) *δ* 7.43-7.37 (m, 1H), 7.25 – 7.17 (m, 3H), 6.87 (d, *J* = 15.2 Hz, 1H), 6.74 (d, *J* = 15.2 Hz, 1H), 4.16 (q, *J* = 7.1 Hz, 2H), 3.90 – 3.77 (m, 2H), 1.25 (t, *J* = 7.1 Hz, 3H), 1.15 (t, *J* = 7.1 Hz, 3H). ESI-HRMS [M+H]^+^ calcd for C_14_H_17_FNO_3_: 266.1187, found: 266.1185.

*(E)-N-ethyl-N-(4-fluorophenyl)but-2-enamide* (**JM13**)

792 mg, 95.5 % yield. **^1^H NMR** (600 MHz, CDCl3) *δ* 7.15 – 7.09 (m, 4H), 6.92 (dq, J = 14.0, 6.8 Hz, 1H), 5.62 (d, J = 14.9 Hz, 1H), 3.78 (q, J = 7.1 Hz, 2H), 1.73 (d, J = 6.8 Hz, 3H), 1.13 (t, J = 7.1 Hz, 3H). ESI-HRMS [M+H]^+^ calcd for C_12_H_15_FNO: 208.1132, found: 208.1132.

*Ethyl (E)-4-(ethyl(4-fluorophenyl)amino)-4-oxobut-2-enoate* (**JM15**)

992 mg, 93.5 % yield. **^1^H NMR** (600 MHz, CDCl_3_) *δ* 7.13 (d, *J* = 6.4 Hz, 4H), 6.84 (d, *J* = 15.3 Hz, 1H), 6.74 (d, *J* = 15.3 Hz, 1H), 4.17 (q, *J* = 7.1 Hz, 2H), 3.83 (q, *J* = 7.1 Hz, 2H), 1.25 (t, *J* = 7.1 Hz, 3H), 1.15 (t, *J* = 7.1 Hz, 3H). ESI-HRMS [M+Na]^+^ calcd for C_14_H_16_FNO_3_Na: 288.1006, found: 288.1010.

*General procedures for the synthesis of* **JM06-JM09, JM11, JM14**

To a solution of **JM10, JM12, JM15, 9-11** (2.0 mmol) in methanol (5 mL) was added 1M NaOH (5 mL). The resulting mixture was stirred at room temperature for 2 h. Then the methanol was removed under reduced pressure, and the residue was acidified to pH = 2 or below with HCl (1M). Then the solution was extracted with ethyl acetate and the combined organic solvents were dried with sodium sulfate and concentrated *in vacuo*. The crude compound was purified by silica gel column chromatography.

*(E)-3-(N-ethylbut-2-enamido)-4-methylbenzoic acid* (**JM06**)

485 mg, 98.0 % yield. **^1^H NMR** (600 MHz, CDCl_3_) *δ* 10.91 (s, 1H), 8.04 (dd, *J* = 8.0, 1.3 Hz, 1H), 7.86 (d, *J* = 1.3 Hz, 1H), 7.43 (d, *J* = 8.0 Hz, 1H), 7.01 (dq, *J* = 13.9, 6.9 Hz, 1H), 5.50 (dd, *J* = 15.0, 1.4 Hz, 1H), 4.11 (dq, *J* = 14.2, 7.2 Hz, 1H), 3.45 (dq, *J* = 14.2, 7.2 Hz, 1H), 2.29 (s, 3H), 1.72 (dd, *J* = 6.9, 1.2 Hz, 3H), 1.18 (t, *J* = 7.2 Hz, 3H). ESI-HRMS [M+H]^+^ calcd for C_14_H_18_NO_3_: 248.1281, found: 248.1281.

*(E)-4-(N-ethylbut-2-enamido)-3-methylbenzoic acid* (**JM07**)

478 mg, 97.0 % yield. **^1^H NMR** (400 MHz, CDCl_3_) *δ* 8.09 (d, *J* = 1.3 Hz, 1H), 8.02 (d, *J* = 8.0 Hz, 1H), 7.22 (d, *J* = 8.1 Hz, 1H), 7.00 (dq, *J* = 13.8, 6.8 Hz, 1H), 5.49 (d, *J* = 14.9 Hz, 1H), 4.09 (dq, *J* = 14.0, 7.1 Hz, 1H), 3.46 (dq, *J* = 13.9, 7.0 Hz, 1H), 2.28 (s, 3H), 1.72 (d, *J* = 6.8 Hz, 3H), 1.17 (t, *J* = 7.1 Hz, 3H). ESI-HRMS [M+H]^+^ calcd for C_13_H_18_NO: 248.1281, found: 248.1281.

*(E)-4-(ethyl(o-tolyl)amino)-4-oxobut-2-enoic acid* (**JM08**)

460 mg, 98.5 % yield. **^1^H NMR** (600 MHz, CDCl_3_) *δ* 7.33 – 7.29 (m, 2H), 7.27 – 7.23 (m, 1H), 7.07 (d, *J* = 7.6 Hz, 1H), 6.83 (d, *J* = 15.3 Hz, 1H), 6.65 (d, *J* = 15.3 Hz, 1H), 4.14 – 4.06 (m, 1H), 3.47 – 3.40 (m, 1H), 2.18 (s, 3H), 1.16 (t, *J* = 7.2 Hz, 3H). ESI-HRMS [M+H]^+^ calcd for C_13_H_16_NO_3_: 234.1125, found: 234.1122.

*(E)-4-(ethyl(p-tolyl)amino)-4-oxobut-2-enoic acid* (**JM09**)

464 mg, 99.5 % yield. **^1^H NMR** (600 MHz, CDCl_3_) *δ* 7.22 (d, *J* = 7.7 Hz, 2H), 7.00 (d, *J* = 7.6 Hz, 2H), 6.80 (s, 2H), 3.83 (q, *J* = 7.1 Hz, 2H), 2.39 (s, 3H), 1.14 (t, *J* = 7.1 Hz, 3H). **^13^C NMR** (150 MHz, CDCl_3_) *δ* 169.96, 163.42, 138.49, 138.02, 136.52, 130.48, 129.85, 127.75, 44.79, 21.12, 12.75. ESI-HRMS [M+H]^+^ calcd for C_13_H_16_NO_3_^+^: 234.1125, found: 234.1129.

*(E)-4-(ethyl(2-fluorophenyl)amino)-4-oxobut-2-enoic acid* (**JM11**)

464 mg, 97.8 % yield. **^1^H NMR** (600 MHz, CDCl_3_) *δ* 7.43 – 7.38 (m, 1H), 7.24-7.16 (m, 3H), 6.83 (d, *J* = 15.2 Hz, 1H), 6.77 (d, *J* = 15.2 Hz, 1H), 3.89 – 3.76 (m, 2H), 1.15 (t, *J* = 7.2 Hz, 3H). ESI-HRMS [M+H]^+^ calcd for C_12_H_13_FNO_3_: 238.0874, found: 238.0874.

*(E)-4-(ethyl(4-fluorophenyl)amino)-4-oxobut-2-enoic acid* (**JM14**)

471 mg, 99.2 % yield. **^1^H NMR** (600 MHz, CDCl_3_) *δ* 7.13 (d, *J* = 6.2 Hz, 4H), 6.79 (q, *J* = 15.3 Hz, 2H), 3.83 (q, *J* = 6.9 Hz, 2H), 1.15 (t, *J* = 7.0 Hz, 3H). ESI-HRMS [M-H]^-^ calcd for C_12_H_11_FNO_3_^-^: 236.0728, found: 236.0724.

## Acknowledgements

This study was supported by the National Natural Science Foundation of China [22037002, 81772689], the Program for Professor of Special Appointment [Eastern Scholar TP2018025] at Shanghai Institutions of Higher Learning, Sponsored by Natural Science Foundation of Shanghai [21ZR1416700], the Innovation Program of Shanghai Municipal Education Commission [2021-01-07-00-02-E00104], and the Chinese Special Fund for State Key Laboratory of Bioreactor Engineering [2060204].

## Author Contributions

Research design: KTB, ZLH and JL. Data collection, analysis, and interpretation: KTB, JLF, WWL, ZFM, TYS, ZZS. Preparation of figure composites: KTB, ZLH and JL. Manuscript writing: KTB, ZLH and JL. All authors provided feedback and edits to the manuscript text and approved the final version of the manuscript.

## Data Availability Statement

The data used to support the findings of this study are provided as figure source data.

---

**Figure S1.**
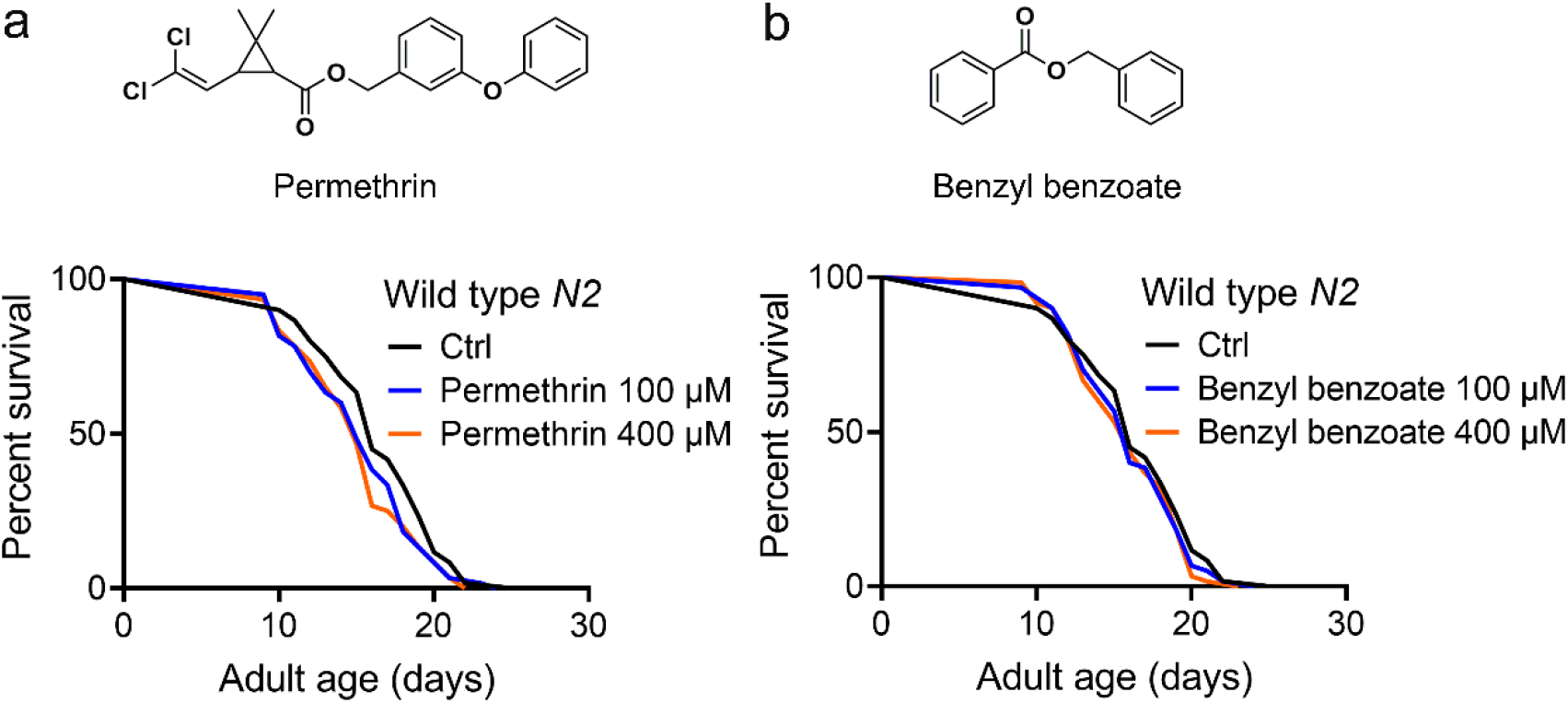
The anti-scabies drugs permethrin and benzyl benzoate failed to extend the lifespan in *C. elegans*. (**a**) Permethrin failed to extend the lifespan of *C. elegans*. (**b**) Benzyl benzoate failed to extend the lifespan of *C. elegans*. (**a**-**b**) Data were compared using the Log-rank test and statistics have been mentioned in Table S1 Experimental group 3. * *P* < 0.05.

**Figure S2.**
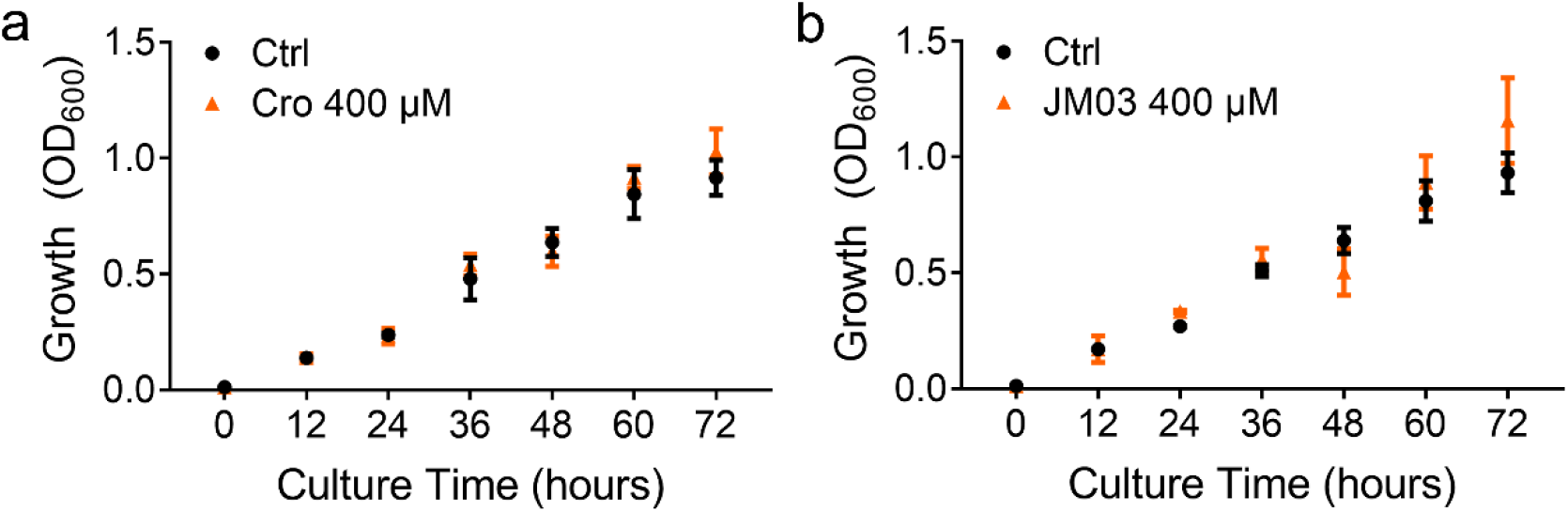
Crotamiton and JM03 did not reduce the bacterial growth at 400 μM concentration. (**a**) Crotamiton did not reduce the bacterial growth at 400 μM concentration. (**b**) JM03 did not reduce the bacterial growth at 400 μM concentration. Data represented as the mean ± SD, and comparisons were made using Student t-test. The graphics represent a compilation of at least 3 independent experiments. * *P* < 0.05.

**Figure S3.**
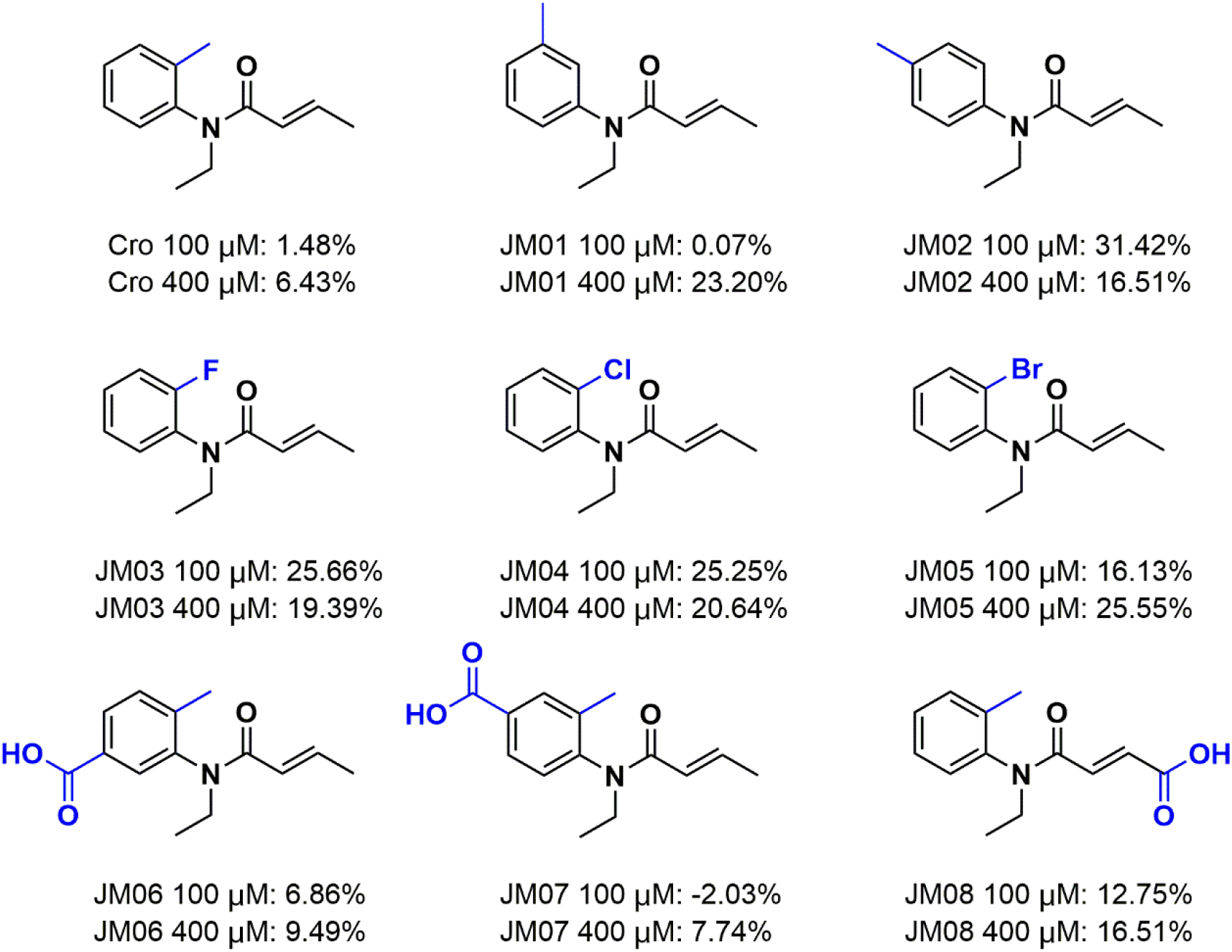
The structures and mean percentage of lifespan extension by crotamiton derivatives.

**Figure S4.**
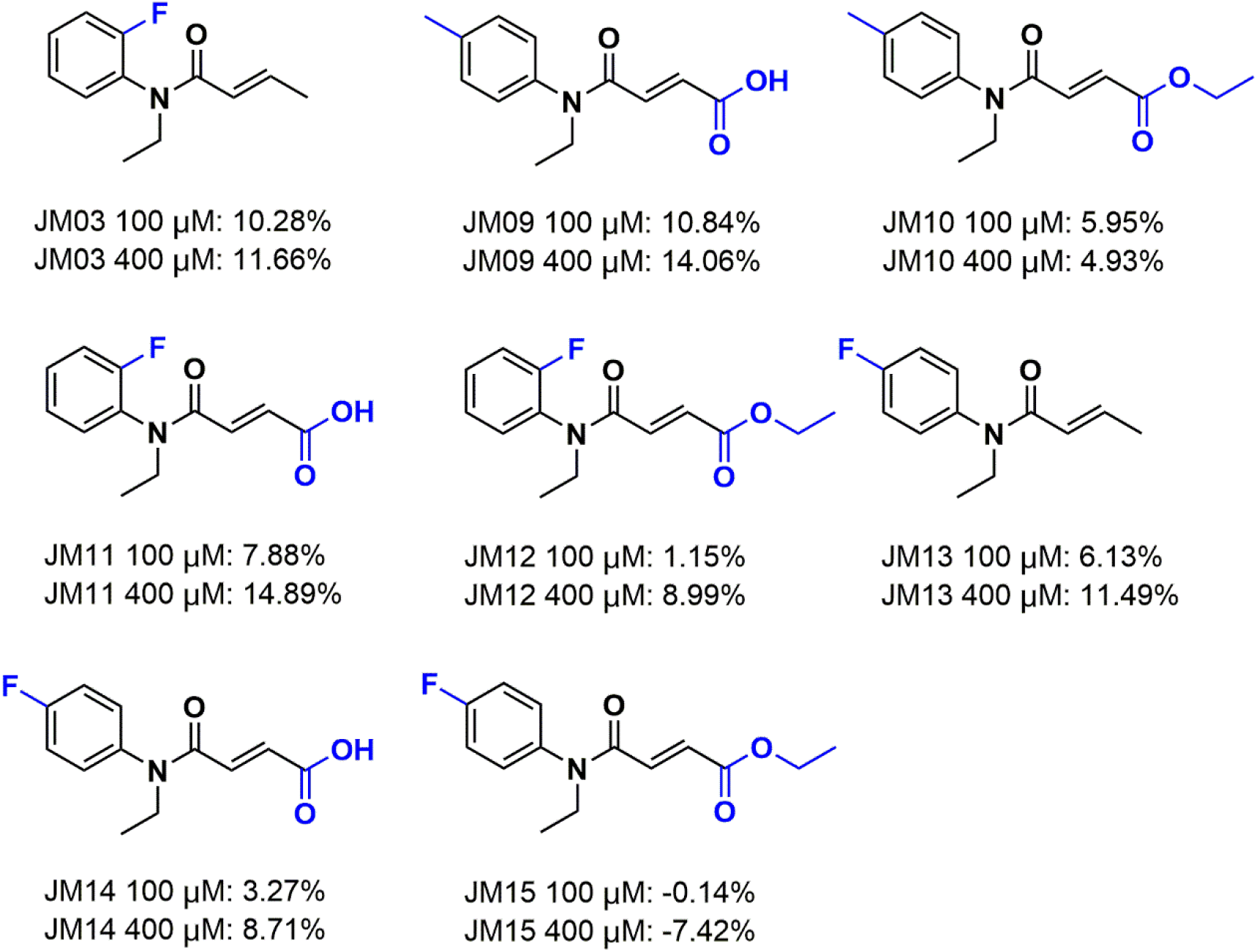
The structures and mean percentage of lifespan extension by crotamiton derivatives.

**Figure S5.**
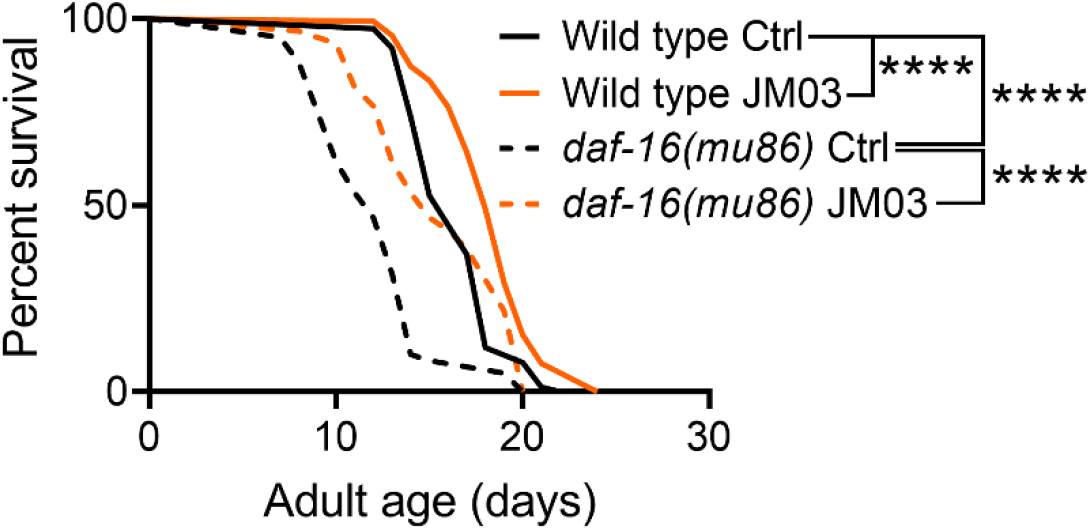
JM03 treatment extended the lifespan of *daf-16(mu86)* mutants. Data are compared using the Log-rank test and statistics have been mentioned in Table S2 Experimental group 2. The graphics represent a compilation of at least 3 independent experiments. \*\*\*\**P*< 0.0001.

